## Supplementary Information for "Inheritable cell-states shape drug-persister correlations and population dynamics in cancer cells"

**4 Euler-Lotka equation connecting single cell kinetics to the population growth rate in the absence of drugs. 7**

**5 Euler-Lotka equation for drug treated cells. 8**

**6 Initial age distribution inference using MLE 9**

**7 Competing risks for exponentially distributed waiting times: an analytical approach. 10**

**8 Competing risks for non-exponential waiting time distributions: Markov Chain Monte Carlo analysis. 13**

**9 Models of cell fate decisions 17**

**10 Simulation of Model *M3* and obtaining the best parameters. 19**

**11 Barcoding simulations** **20**

**12 Calculation of Barcode abundance and diversity** **21**

**13 Luria-Delbruck Analysis** **21**

### 1 Details of the HCT116 and U2OS datasets.

We analyzed two datasets – time lapse imaging data for HCT116 [1] and U2OS cells [2]. In the HCT116 dataset, both daughters of a mother cell were tracked, while in the U2OS dataset only one randomly chosen daughter was tracked. The data for HCT116 cells had 83 ancestor cells which gave rise to distinct lineage trees. However, we analyzed only 65 of these trees. The remaining 18 were discarded as they either didn't divide at all or had many missing cells in the lineage. The 65 cells at the beginning of the experiment expanded to 275 cells in the two days of drug-free expansion. HCT116 cells were exposed to a single concentration of cisplatin approximately near its IC50 value [1]. U2OS cells on the other hand were treated with four concentrations of cisplatin, 0, 7, 10 and 13 $\mu$M corresponding to Control, Low, Medium and High labels [2], which started with 300, 247, 262 and 316 cells respectively. After removing lineages where the ancestor cell did not divide at all, we were left with 289, 232, 240 and 296 lineages. Besides the cells that did not divide, cells that died before exposure to cisplatin were also excluded from the analysis. Since only one of the daughters of dividing U2OS cells were randomly chosen for tracking, the number of cells at the timepoint of cisplatin addition was equal to that at the start of the experiment. Besides measurements of IMT and AT of single cells, these datasets allow simultaneous extraction of the population dynamics by counting the number of surviving cells in each time frame. Of the 275 HCT116 cells at the start of the three day cisplatin treatment period, 182 died while the remaining 93 either divided during the treatment period or survived without dividing till the end of experiment. Furthermore, the HCT116 dataset also provides information on lineage relationships between the cells.

### 2 Biphasic decay of HCT116 cells and slope estimation using piece-wise linear regression.

From the single cell lineage tracking data for HCT116 cells, we counted the total number of cells in every 30 mins of the biphasic curve post cisplatin exposure and fit the data to a piece-wise linear regression model. The fit was performed using the 'segmented' function of 'segmented' package in R. The function took the data where the exponential nature of decay was observed, i.e. from 80 to 120 hours of the experiment, discarding the transient phase immediately after cisplatin treatment.

The equation for the above fit and the fit statistics are provided below:

$$\log(N(t)) = \gamma_1 * t + (t - \beta) * \gamma_2 + \mathrm{c} \tag{1}$$

Where $N$ is the number of cells and $t$ is time in hours. $\gamma_1$ and $\gamma_2$ are the slopes of the fast and slow decay respectively. $\beta$ is the break-point or the point of transition in hours, where the fast and slow decay slopes intersect.

The parameters after the fit for the population decay of HCT116 cells are $\gamma_1 = -0.0286\, h^{-1}$, $\gamma_2 = -0.0032\, h^{-1}$ and $\beta = 102\, h$.

### 3 Estimating Intermitotic Time (IMT) distributions of pre-cisplatin HCT116 and U2OS cells.

#### 3.1 Inferring the parameters of the best-fitting IMT distribution

Previous studies have shown that the IMT distribution from drug naive cells is best fit with an Exponentially Modified Gaussian (EMG) probability density function [1]. This model has 3 parameters, $\mu$, $\sigma$ and $\lambda$, the first two parameterizing the Gaussian and the last parameterizing the rate of the exponential. Maximum Likelihood estimation was used to infer the parameters of the EMG. Let $t_i$ be the IMT value of the $i^{th}$ cell (only cells that underwent a complete division

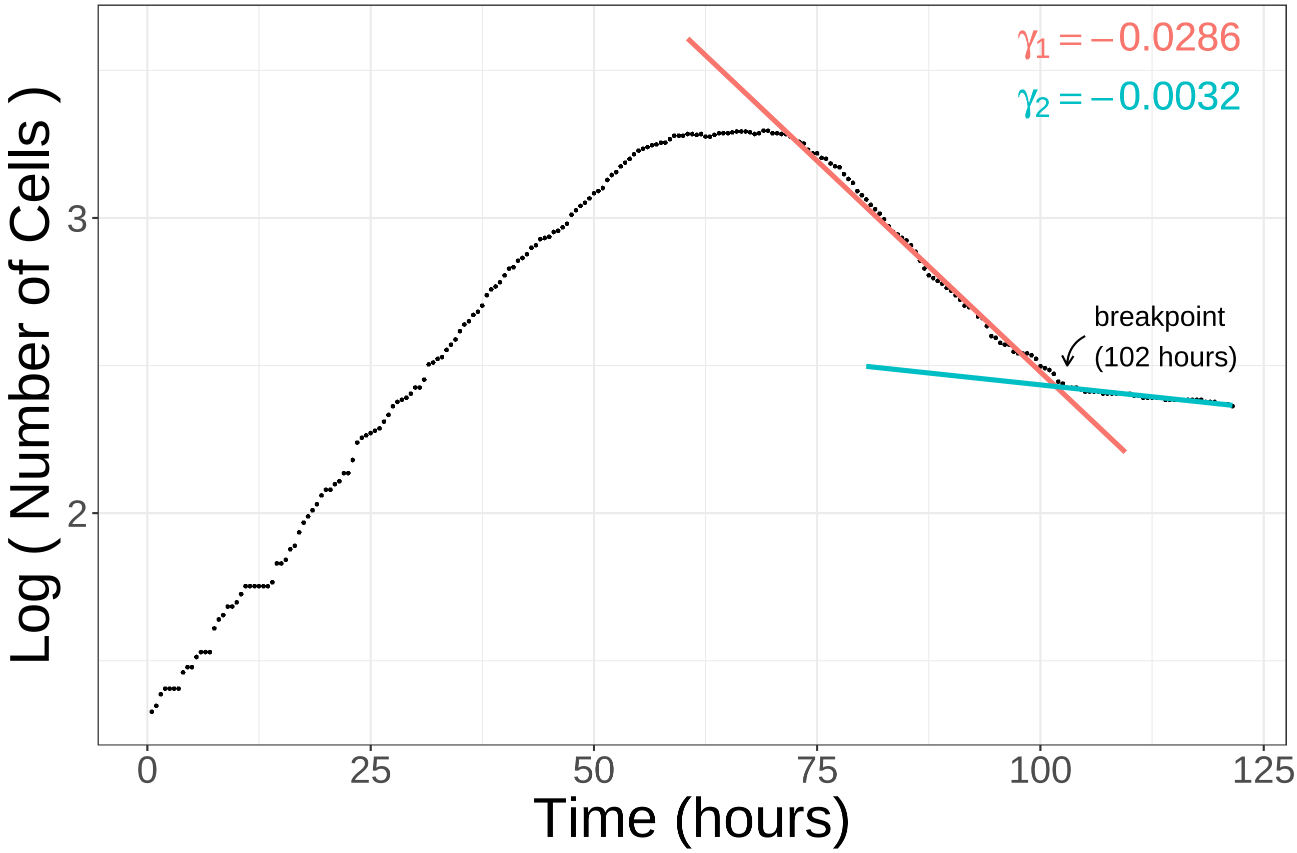

Supplementary Figure 1: Bi-phasic population decay curve for HCT116 cells. The bi-phasic decay post cisplatin treatment shows two distinct slopes, referred to in the legend as $\gamma_1$ and $\gamma_2$. The break-point $\beta$ is the point of intersection of the two slopes. The second slow decay arises from the persister population.

are considered) and $f(t_i; \boldsymbol{\Theta}_{\mathrm{m}}^{\mathrm{bef}})$ be its corresponding probability density with parameter vector $\boldsymbol{\Theta}_{\mathrm{m}}^{\mathrm{bef}}$ denoting the parameters of the EMG distribution. Then the likelihood $L(\boldsymbol{\Theta}_{\mathrm{m}}^{\mathrm{bef}}|\mathbf{t})$ can be written as:

$$L(\boldsymbol{\Theta}_{\mathrm{m}}^{\mathrm{bef}}|\mathbf{t}) = \Pi_i f_i(t_i; \boldsymbol{\Theta}_{\mathrm{m}}^{\mathrm{bef}}) \quad (2)$$

where, $i$ iterates over the IMT values of all the drug naive cells dividing before the addition of cisplatin.

To find the Maximum Likelihood Estimate of the EMG parameters, this likelihood function was maximised using the 'emg.mle' command from the 'emg' package in R using the initial parameters $\mu = 80$ h, $\sigma = 10^{-2}$ h and $\lambda = 10^{-2}\,\mathrm{h}^{-1}$ . Lower bounds for $\mu = 1$ h, $\sigma = 10^{-4}$ h and $\lambda = 10^{-4}\,\mathrm{h}^{-1}$ were also provided to 'emg.mle', to prevent the parameters from becoming 0 or taking negative values.

The EMG parameter values thus obtained for HCT116 were $\mu = 13.97\,\mathrm{h}$, $\sigma = 1.11\,\mathrm{h}$ and

$\lambda = 0.33\,\mathrm{h}^{-1}$; while for U2OS, the parameters are provided in Table 1 below:

| Conc. | $\mu$ (h) | $\sigma$ (h) | $\lambda$ ($\mathrm{h}^{-1}$) |
|---|---|---|---|
| High | 24.32 | 1.54 | 0.32 |
| Medium | 23.92 | 2.40 | 0.4 |
| Low | 22.56 | 2.16 | 0.39 |

Table 1: EMG parameters for IMT distributions of drug-naive U2OS cells. The three concentrations High, Medium and Low denote the three experimental conditions. Since these parameters describe IMT distributions before cisplatin addition, they are nearly the same across the experimental conditions.

##### 3.2 Segregating lineages of persister and sensitive cells in the HCT116 dataset.

There was a clear biphasic exponential decay in the HCT116 data post cisplatin, with cells surviving till the end of experiment contributing to the second slow exponential decay. These surviving cells were identified as persisters. To check whether the ancestors of the persister and sensitive cells (cells that died post cisplatin) displayed any differences in cycling rates, we extracted all the lineages giving rise to the two phenotypes (survival versus death) and compared their division times. Since many ancestors in the lineages of either phenotypes were common, we resorted to the following analysis. First, we found all the ancestor cells in the lineages that led to a specific phenotype from the lineage tree, say persistence. Next, we compared these ancestors amongst the two phenotypes (persistence versus death) and extracted only the ones unique to the persistence phenotype. Similarly the ancestors unique to the sensitive phenotype were extracted. The IMT distributions of these ancestor cells of the two phenotypes were then plotted and compared (Fig. 1c of the main text).

##### 3.3 Mixture model for the U2OS dataset.

The single-cell timelapse data for U2OS had only one of the daughters tracked after every division event. The data thus provided us only with forward lineages. Since we couldn't compare the IMTs of the persister and sensitive cells' lineages separately, for any given lineage tree, we approached the problem by fitting a mixture model to the IMT distribution and comparing the fit with that of single EMG using the Akaike Information Criterion (AIC). Since the experiment in U2OS was carried out on three different concentrations of cisplatin, we could check for consistency of our results by repeating this procedure on three separate drug naive cell populations. We observed that for all the three cases, the single EMG model fit the IMT distribution better than the mixture model. For single EMG model fit, we used Maximum Likelihood Estimate (MLE) as described before. For the mixture model, however, a custom likelihood function in R was written and its parameters estimated using a Nonlinear Least Squares fit provided by 'nls_multstart' function from the 'nls.multstart' package in R. The mixture model assumes that the IMT values can be fit with two EMG distributions having distinct parameter values. Along with the parameters of the EMGs it also calculates the weight of each EMG distribution. Table 2 provides the weight of one distribution and the best fit parameter values of the two EMGs.

| **Conc.** | $w_1$ | $\mu_1$ | $\sigma_1$ | $\lambda_1$ | $\mu_2$ | $\sigma_2$ | $\lambda_2$ |
|---|---|---|---|---|---|---|---|
| High | 0.85 | 24.37 | 1.65 | 0.39 | 30.46 | 5.11 | 7.08 |
| Medium | 0.81 | 24.99 | 2.37 | 0.64 | 19.47 | 1.22 | 0.15 |
| Low | 0.94 | 22.57 | 1.91 | 0.33 | 20.05 | 1.05 | 31.83 |

Table 2: Parameters of the mixture model of 2 EMGs, after fitting to the pre-cisplatin U2OS datasets.

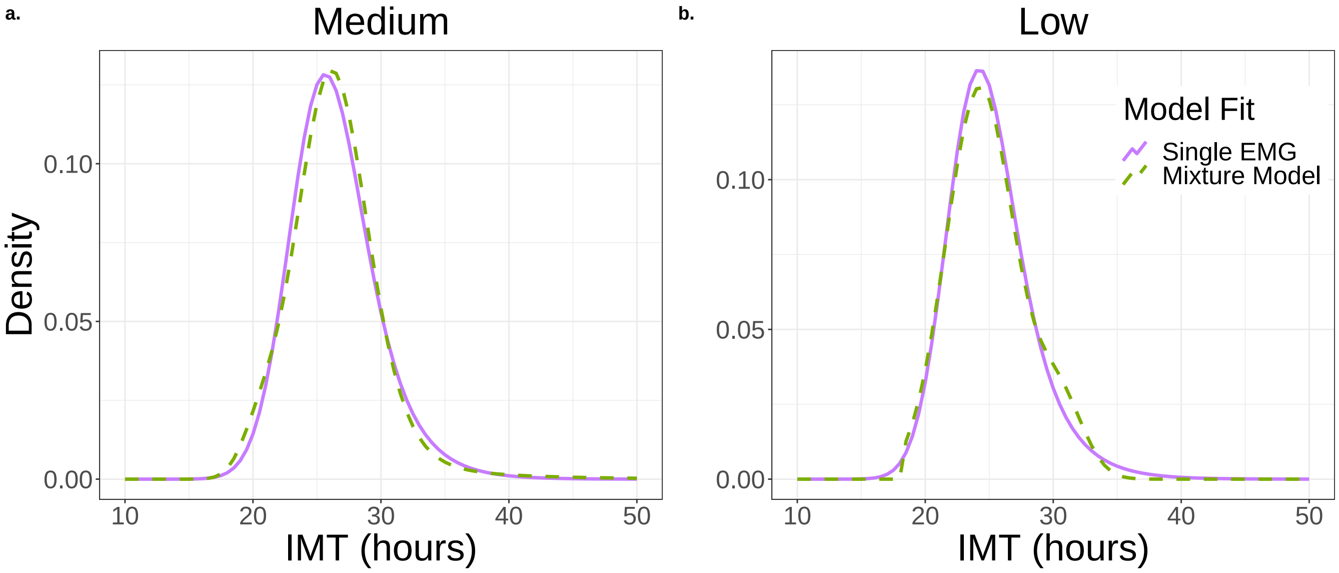

Supplementary Figure 2: Single EMG fits to pre-cisplatin U2OS data perform better than mixtures of EMGs. (a) Data from medium cisplatin concentration. (b) Data from low cisplatin concentration. The results for the High cisplatin concentration are shown in Main Text Figure 1f.

AIC values of the two models are shown in Table 3 below:

| Model | High | Medium | Low |
|---|---|---|---|
| Single EMG | 862.64 | 816.08 | 954.24 |
| Mixture EMG | 873.74 | 819.79 | 962.16 |

Table 3: AIC values to compare fits of single EMG versus mixture of EMG models, for the pre-cisplatin U2OS datasets. In each experimental condition High, Medium and Low, the single EMG model has a lower AIC value than the mixture of EMGs, demonstrating that the single EMG explains the data better in all cases.

### 4 Euler-Lotka equation connecting single cell kinetics to the population growth rate in the absence of drugs.

In an exponentially growing population of cells starting with $N(0)$ number of cells, the average size of the population at time $t$, $N(t)$, is given by

$$N(t) = N(0)e^{\gamma t} \tag{3}$$

where $\gamma$ is the population growth rate.

If in one lineage the first ancestor cell divides after time $\tau$ to give rise to two daughter cells, then the cell population within that lineage at time $t$ can be thought to have arisen from the descendants of the two daughters in time $t - \tau$. Since $\tau$ is a random number drawn from the IMT distribution $f(\tau)$, the average number of cells across all $N(0)$ lineages at time $t$, $N(t)$, will be given by the following equation:

$$N(t) = 2N(0)\langle e^{\gamma(t-\tau)}\rangle, \tag{4}$$

where the angular brackets denote an average over the IMT distribution $f(\tau)$. Equation 4 can be rewritten as:

$$N(t) = 2N(0)\int_0^\infty e^{\gamma(t-\tau)}f(\tau)d\tau. \tag{5}$$

Equating equations 3 and 5, we obtain the Euler Lotka equation [3, 4]:

$$1 = 2\int_0^\infty e^{-\gamma\tau} f(\tau) d\tau. \tag{6}$$

The Euler-Lotka equation is an integral equation connecting the single cell IMT distribution $f(\tau)$ to the population growth rate $\gamma$ in the absence of any cell death. We use the experimentally measured $f(\tau)$ and solve the Euler-Lotka equation numerically to obtain the growth rate $\gamma$.

### 5 Euler-Lotka equation for drug treated cells.

In the presence of cell death (which may occur due to drug treatment), a standard approach to estimating the population growth (or decay) rate from the single cell kinetics is using the Von-Foerster equation [5]. This equation relates the rate of change of cell population of age $a$ at time $t$ to the age dependent hazards of division and death. The age of a cell is defined as the time interval that the cell survives without dividing or dying when measured from the time of its birth.

Let $n(t, a)$ be the number of cells of age $a$ ($\forall\, a \geq 0$) at time $t$ . Then,

$$\frac{\partial n(t,a)}{\partial t} + \frac{\partial n(t,a)}{\partial a} = -(\mu(a) + b(a))n(t,a), \tag{7}$$

where, $b(a)$ and $\mu(a)$ are the age-dependent birth and death hazards. For the newborn cells i.e. n(t,0),

$$n(t,0) = \int_0^\infty b(a)n(t,a)dt. \tag{8}$$

In the long time limit, the number of cells grow exponentially and so by using similarity method of solving partial differential equations (PDEs), we would obtain

$$n(t,a) = e^{\gamma t} r(a), \tag{9}$$

where, $\gamma$ is the growth (or decay) rate and $r(a)$ is the age distribution of the cell population in the long time limit. Substituting 9 in 7 we obtain an ordinary differential equation (ODE) for the steady state age distribution $r(a)$,

$$\frac{dr}{da} = -(b(a) + \mu(a) + \gamma)r(a). \tag{10}$$

Simplifying it further,

$$r(a) = r(0)e^{-\gamma a - \int_0^a (b(s)+\mu(s))ds}. \tag{11}$$

The exponentially growing cell population of age $a$ at time $t$ is given by

$$n(t, a) = e^{\gamma t} r(0) e^{\gamma a - \int_0^a (b(s)+\mu(s))ds}. \tag{12}$$

Now, substituting 12 into 8 and simplifying we get the Euler-Lotka equation for cells in the presence of a drug [5]

$$1 = 2\int_0^\infty b(a)e^{-\gamma a}e^{-\int_0^a (b(s)+\mu(s))ds}da. \tag{13}$$

.

As in the previous case of the Euler-Lotka equation in the absence of cell death, the Euler-Lotka equation in the presence of cell death is also an integral equation. We use the experimentally measured (or the MCMC inferred; see Section 8 below) IMT and AT distributions post drug to convert them into the corresponding hazards $b(s)$ and $\mu(s)$ using Eq. 30. These hazards are then used to solve the Euler-Lotka equation numerically to solve for the population decay rate $\gamma$.

### 6 Initial age distribution inference using MLE

The predictions of population growth rate from Euler-Lotka equations are dependent on the initial age distribution of the cells. Since cells at the start of the experiment are provided fresh media, they tend to undergo synchronisation. This effectively leads to delays in the times to division of the cells. These delays manifest in the average population time curve as step-like patterns called 'mitotic-waves' which dampen over long time observations [6]. Since the age distribution of the cells at the start of the experiment was unknown, we inferred this distribution using the following simple likelihood maximization procedure:

Let the IMT of the cell be $t_{\mathrm{IMT}}$. This is assumed to be distributed as the measured IMT of the drug naive cells in the experiment. Let $\tau_a$ be the age of the cell at the start of the experiment and $\tau_D$ be its time to division. Then, the probability density that this cell divides at $t_{IMT}$ given that it has survived upto time $\tau_a$ is,

$$f(t_{\mathrm{IMT}} = \tau_a + \tau_D | t_{\mathrm{IMT}} > \tau_a) = \frac{f(t_{\mathrm{IMT}} = \tau_a + \tau_D, t_{\mathrm{IMT}} > \tau_a)}{f(t_{\mathrm{IMT}} > \tau_a)} \tag{14}$$

$$f(t_{\mathrm{IMT}} = \tau_a + \tau_D | t_{\mathrm{IMT}} > \tau_a) = \frac{f(t_{\mathrm{IMT}} = \tau_a + \tau_D)}{f(t_{\mathrm{IMT}} > \tau_a)} \tag{15}$$

where,

$$t_{\mathrm{IMT}} = \tau_a + \tau_D \tag{16}$$

The most likely age of the cell was then determined as the age at which this conditional probability peaked.

### 7 Competing risks for exponentially distributed waiting times: an analytical approach.

Here we provide an analytical calculation to demonstrate how the competing risks effect precludes estimation of the underlying IMT and AT distribution parameters directly from measurements of cell division (mitosis) and death (apoptosis) time. We also show how the underlying distributions can accurately be inferred by writing down the Likelihood function for the data and inferring the real parameters. Here we assume all distributions are exponential, to allow analytical solutions. The more complex case of non-exponential IMT and AT distributions is treated in the next section.

For a cell, let the probability densities of waiting times to mitosis and apoptosis be given by $f_m(t)$ (Supplementary Figure 3a) and $f_a(t)$ (Supplementary Figure 3d). Assuming these to be exponential, their probability densities can be written as

$$f_m(t) = \lambda_m e^{-\lambda_m t} \quad (17) \qquad f_a(t) = \lambda_a e^{-\lambda_a t} \quad (18)$$

Let 'E' be an event that occurs at time 't'. In the current scenario event E is a discrete random variable that can either be apoptosis or mitosis. The time $t$ however is a continuous random variable defining the minimum time required for any of the events to occur. The probabilities of the events and probability density if the time to the event are:

$$P(\mathrm{E} = \text{Apoptosis}) = \frac{\lambda_a}{\lambda_a + \lambda_m} \quad (19) \qquad P(\mathrm{E} = \text{Mitosis}) = \frac{\lambda_m}{\lambda_a + \lambda_m} \quad (20)$$

$$f_t(t_E) = (\lambda_a + \lambda_m) e^{-(\lambda_a + \lambda_m) t_E} \quad (21)$$

Since a cell cannot undergo both mitosis and apoptosis simultaneously, the events are mutually exclusive. However, these events are not independent. For this reason, in the presence of the drug, the probability of an event is effected by the competing risk from the other. The modified probability densities are calculated as a product of probability of an event with the probability density of the time to an event. The modified densities are shown below for events of mitosis and apoptosis respectively.

$$g_m(t) = \lambda_m e^{-(\lambda_m + \lambda_a) t} \quad (22) \qquad g_a(t) = \lambda_a e^{-(\lambda_m + \lambda_a) t} \quad (23)$$

By tracking single cells, both the sequence of events and the times to those events can be obtained. If $E_i$ be the $i^{th}$ event in the sequence of events and $t_i$, the waiting time to the $i^{th}$ event, finding rates of the waiting time distributions to the events becomes an inverse problem. If there are $N_m$ observations of mitosis and $N_a$ of apoptosis, the likelihood of observing this data given the exponential distributions is given by

$$L(\lambda_m, \lambda_a) = \lambda_m^{N_m} \lambda_a^{N_a} e^{-(\lambda_m + \lambda_a) \sum_1^{N_m + N_a} t_i}. \quad (24)$$

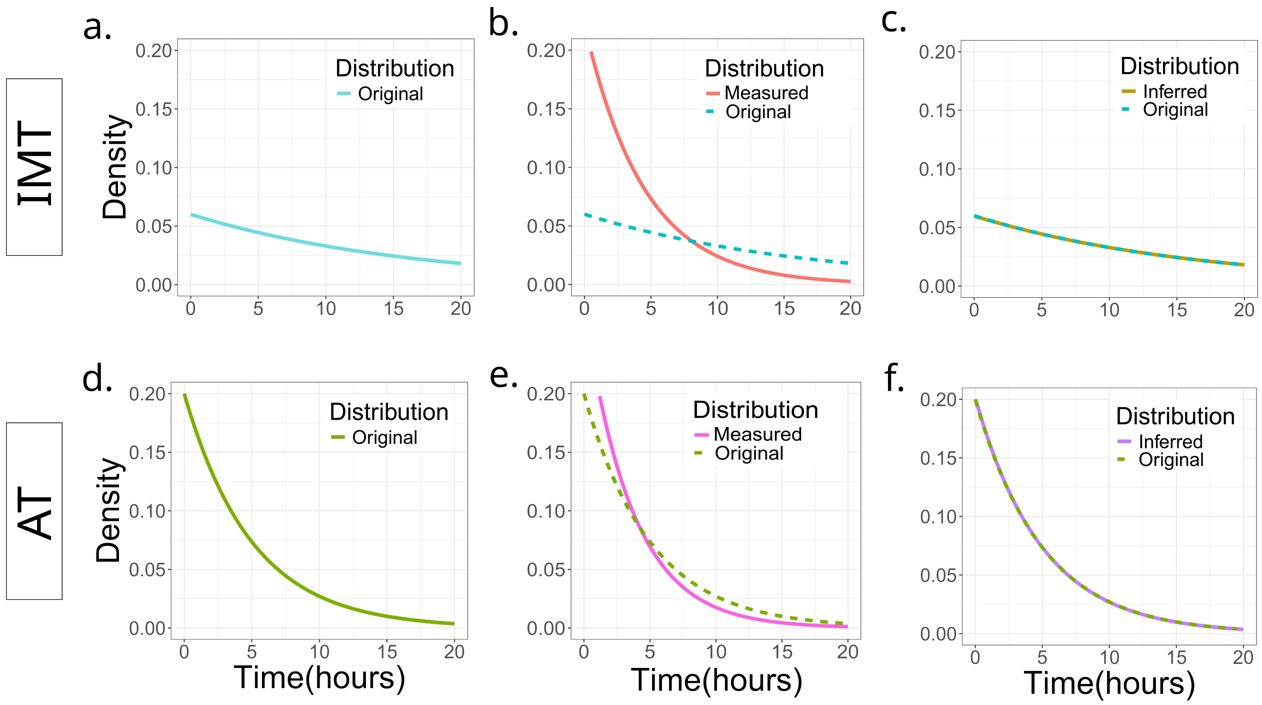

Supplementary Figure 3: Inferring the original exponential distributions of IMT and AT in the presence of competing risks using an analytical approach. (a)-(c) are exponential IMT distributions; (d)-(f) are exponential AT distributions. (a) and (d) show the underlying original distributions that were used in the simulations. (a) The original distribution of IMT used in the simulations was an exponential distribution with hazard $\lambda_m^{\text{orig}} = 0.06h^{-1}$. (b) 'Measured' distribution shows the exponential fit to the measured IMT distribution. IMT values from cells tracked in the simulation were extracted and their density distribution was fit to an exponential using Maximum Likelihood Estimate (MLE). The hazard of 'Measured' was $\lambda_m^{\text{meas}} = 0.22h^{-1}$ . The 'Measured' shows a reduced right tail compared to the 'Original' distribution (dotted). (c) 'Inferred' shows the exponential distribution with hazard inferred using equation 25. When compared to the 'Original' distribution the two match with an error of less than 1%. The hazard of the inferred distribution was $\lambda_m^{\text{inf}} = 0.06h^{-1}$. (d) The original distribution of AT used in the simulations was an exponential distribution with hazard $\lambda_a^{\text{orig}} = 0.20h^{-1}$. (e) 'Measured' AT distribution obtained by fitting an exponential to the AT obtained from the simulation shows a reduced right tail compared to the 'Original' distribution. AT values from the simulation were extracted and their density distribution was fit to an exponential using MLE. The hazard of the 'Measured' AT was $\lambda_a^{meas} = 0.27h^{-1}$. (f) 'Inferred' shows the exponential distribution with hazard inferred using equation 26. When compared to the 'Original' distribution the two match with an error of less than 1%. The hazard of the 'Inferred' distribution was $\lambda_a^{\text{inf}} = 0.20h^{-1}$.

The real underlying rates of inter-mitosis and apoptosis times can then be obtained by maximising the likelihood with respect to $\lambda_m$ and $\lambda_a$. Since here the likelihood is differentiable, the maxima can be analytically calculated using method of derivatives. The inferred rates can then be written as:

$$\lambda_m^{\text{inf}} = \frac{N_m}{\sum_1^{N_m+N_a} t_i} \qquad (25) \qquad\qquad \lambda_a^{\text{inf}} = \frac{N_a}{\sum_1^{N_m+N_a} t_i} \qquad (26)$$

The inferred rates in Equations 25 and 26 match within an error of 1% with the rates used in the original distribution (Supplementary Figures 3c and f). This highlights the necessity of using inference to correctly establish the underlying distributions when competing risks exist.

### 8 Competing risks for non-exponential waiting time distributions: Markov Chain Monte Carlo analysis.

#### 8.1 Survival analysis

If $T$ is a non-negative continuous random variable, then $f(t)$ denotes the probability density function of $T$. Furthermore, the cumulative distribution function (CDF), $F(t)$ is defined as

$$F(t) = Pr(T \leq t) = \int_0^t f(s)ds. \tag{27}$$

A term often used in survival analysis is survival probability $S(t)$ which denotes the probability that the event has not occurred by time t:

$$S(t) = Pr(T > t) = 1 - F(t) \tag{28}$$

An instantaneous rate of occurrence of the event or hazard $h(t)$, is defined as:

$$h(t) = \lim_{\delta t \to 0} \frac{Prob(t < T \leq t + \delta t | T > t)}{\delta t} \tag{29}$$

Equation 29 for the hazard can then be simplified to be a ratio of probability density and survival probability as shown below:

$$h(t) = \frac{f(t)}{S(t)} \tag{30}$$

This relation between probability density $f(t)$, survival probability $S(t)$ and the equation 28 allows to write the survival probability completely in terms of the hazard as:

$$S(t) = e^{-\int_0^t h(s)ds} \tag{31}$$

#### 8.2 Hazard in the presence of competing risks

Both U2OS and HCT116 cells exposed to the cisplatin were divided into two categories based on their birth time. While representing the time of drug addition as $T_d$, we accounted for the delay in cisplatin effects by assuming a parameter $T_{\text{delay}}$. The cells born before $T_d + T_{\text{delay}}$ and surviving this delay period, experienced a shift in their hazards. These cells transitioned from undergoing just division in drug naive condition to experiencing a competing risk between division and death. The hazard function for straddling cells can be defined as a piece-wise function shown below:

$$h^i_{\text{straddle}}(t) = \begin{cases} h(t; \mathbf{\Theta}^{\text{bef}}_{\text{m}}), & t < T_d + T_{\text{delay}} - T^i_{\text{birth}} \\ h(t; \mathbf{\Theta}^{\text{aft}}_{\text{m}}) + h(t; \mathbf{\Theta}^{\text{aft}}_{\text{a}}), & t \geq T_d + T_{\text{delay}} - T^i_{\text{birth}} \end{cases}, \tag{32}$$

where $h(t; \mathbf{\Theta}^{\text{aft}}_{\text{m}})$ and $h(t; \mathbf{\Theta}^{\text{aft}}_{\text{a}})$ denote the post-cisplatin division and death hazards of a cell respectively. $\mathbf{\Theta}^{\text{bef}}_{\text{m}}$ denotes the parameter vector of the pre-drug IMT distribution while $\mathbf{\Theta}^{\text{aft}}_{\text{m}}$ and $\mathbf{\Theta}^{\text{aft}}_{\text{a}}$ denote the parameter vectors parameterizing the post-drug IMT and AT distributions respectively. The time of cisplatin addition, $T_d$ for HCT116 and U2OS were 49 and 48 hours respectively. The delay time, $T_{\text{delay}}$ is obtained from MCMC sampling of the posterior mentioned below.

#### 8.3 Survival in the presence of competing risks

Cells exposed to cisplatin face a competition in the fates of division and death. In such a competing risks scenario, the fate of a cell is affected by the combined hazards of the two fates. This combined hazard is faced by the cell when the effects of cisplatin kick in, i.e. after $t \geq T_d + T_{\text{delay}}$.

$$S^{\text{strad}}_{\text{Total}}(t) = e^{-\int_0^t h_{\text{straddle}}(s)ds}, \tag{33}$$

where $h_{\text{straddle}}$ is defined in Equation 32. Therefore the survival function for the straddling cells can be defined as follows:

$$S^{\text{strad}}_{\text{Total}}(t) = \begin{cases} e^{-\int_0^{T_d+T_{\text{delay}}} h(s;\mathbf{\Theta}^{\text{bef}}_{\text{m}})ds}, & t < T_d + T_{\text{delay}} \\ e^{-\int_{T_d+T_{\text{delay}}}^{t} (h(s;\mathbf{\Theta}^{\text{aft}}_{\text{m}})+h(s;\mathbf{\Theta}^{\text{aft}}_{\text{a}}))ds} & t \geq T_d + T_{\text{delay}}\ , \end{cases} \tag{34}$$

For cells born after cisplatin, the survival probability is calculated as shown below:

$$S^{\text{aft}}_{\text{Total}}(t; \mathbf{\Theta}^{\text{aft}}_{\text{m}}, \mathbf{\Theta}^{\text{aft}}_{\text{a}}) = e^{-\int_0^t (h(s;\mathbf{\Theta}^{\text{aft}}_{\text{m}})+h(s;\mathbf{\Theta}^{\text{aft}}_{\text{a}}))ds} \tag{35}$$

#### 8.4 Probability of a fate in the presence of competing risks

Given that a cell has survived up to time $t$, the probability density that it will undergo a fate of division (mitosis - m) from a choice between competing fates division (mitosis - m) and death (apoptosis - a) is given by the equation below

$$f(t; \mathbf{\Theta}^{\text{aft}}_{\text{m}}, \mathbf{\Theta}^{\text{aft}}_{\text{a}}) = h(t; \mathbf{\Theta}^{\text{aft}}_{\text{m}}) S_{\text{Total}}(t; \mathbf{\Theta}^{\text{aft}}_{\text{m}}, \mathbf{\Theta}^{\text{aft}}_{\text{a}}) \tag{36}$$

#### 8.5 Quiescent cells

Cells that didn't divide until the end of the experiment, could either be right censored or could have transitioned into quiescent cell types. These cells are not actively proliferating and halt their progression through cell cycle in G0 cell cycle phase. To account for these cells, we assumed a cell's shift to quiescence with probability $q$. Thus if the cell undergoes any event (division or death), then the cell evades going to quiescent state with a probability of $1 - q$.

#### 8.6 MCMC to infer the true post-cisplatin IMT and AT distributions.

To infer the parameters of the underlying distribution of IMT and AT for cisplatin exposed cells we used a likelihood approach coupled with Markov Chain Monte Carlo. The IMT and AT data of cisplatin exposed cells can be divided into 4 categories, viz. IMT and AT for straddling cells; and IMT and AT for cells born after exposure to cisplatin. Certain cells were also observed to stay intact without dividing or dying post exposure to cisplatin. These cells can again be of two types, straddling and non-straddling. Likelihoods for each of these types of IMT and AT values were written separately and then combined to give the full likelihood. A Metropolis Hastings algorithm was used to sample from the posterior. The proposal distribution used was a Gaussian with fixed standard deviation values. 200000 iterations were used to sample the posterior. A burn in of 20000 was kept to minimize the effect of initial values.

The likelihoods used for different categories of the data are provided below:

Likelihood for cells born before $T_{\mathrm{d}}$ which undergo mitosis after $T_{\mathrm{d}}$:

$$L_1(\mathbf{\Theta}_{\mathrm{m}}^{\mathrm{aft}}, \mathbf{\Theta}_{\mathrm{a}}^{\mathrm{aft}}, T_{\mathrm{delay}}, q; \mathbf{t}_{\mathrm{m}}^{\mathrm{strad}}) = \prod_{t_i} (1-q) h(t_i; \mathbf{\Theta}_{\mathrm{m}}^{\mathrm{aft}}) S_{\mathrm{Total}}^{\mathrm{strad}}(t_i; \mathbf{\Theta}_{\mathrm{m}}^{\mathrm{aft}}, \mathbf{\Theta}_{\mathrm{a}}^{\mathrm{aft}}), \forall t_i \in \mathbf{t}_{\mathrm{m}}^{\mathrm{strad}}, \tag{37}$$

where $S_{\mathrm{Total}}^{\mathrm{strad}}$ is given in Equation 34.

Cells born before $T_{\mathrm{d}}$ and undergo apoptosis after $T_{\mathrm{d}}$:

$$L_2(\mathbf{\Theta}_{\mathrm{m}}^{\mathrm{aft}}, \mathbf{\Theta}_{\mathrm{a}}^{\mathrm{aft}}, T_{\mathrm{delay}}, q; \mathbf{t}_{a}^{\mathrm{strad}}) = \prod_{t_i} (1-q) h(t_i; \mathbf{\Theta}_{\mathrm{a}}^{\mathrm{aft}}) S_{\mathrm{Total}}^{\mathrm{strad}}(t_i; \mathbf{\Theta}_{\mathrm{m}}^{\mathrm{aft}}, \mathbf{\Theta}_{\mathrm{a}}^{\mathrm{aft}}), \forall t_i \in \mathbf{t}_{\mathrm{a}}^{\mathrm{strad}}. \tag{38}$$

Cells born before $T_{\mathrm{d}}$ and undergo neither apoptosis nor mitosis after $T_{\mathrm{d}}$:

$$L_3(\mathbf{\Theta}_{\mathrm{m}}^{aft}, \mathbf{\Theta}_{\mathrm{a}}^{aft}, T_{\mathrm{delay}}, q; \mathbf{t}_{\mathrm{stay}}^{\mathrm{strad}}) = \prod_{t_i} (q + (1-q) S_{\mathrm{Total}}^{\mathrm{strad}}(t_i; \mathbf{\Theta}_{\mathrm{m}}^{\mathrm{aft}}, \mathbf{\Theta}_{\mathrm{a}}^{\mathrm{aft}})), \forall t_i \in \mathbf{t}_{\mathrm{stay}}^{\mathrm{strad}}. \tag{39}$$

Cells born after $T_{\mathrm{d}}$ and undergo mitosis after $T_{\mathrm{d}}$:

$$L_4(\mathbf{\Theta}_{\mathrm{m}}^{\mathrm{aft}}, \mathbf{\Theta}_{\mathrm{a}}^{\mathrm{aft}}, T_{\mathrm{delay}}, q; \mathbf{t}_{\mathrm{a}}^{\mathrm{aft}}) = \prod_{t_i} (1-q) h(t_i; \mathbf{\Theta}_{\mathrm{m}}^{\mathrm{aft}}) S_{\mathrm{Total}}^{\mathrm{aft}}(t_i; \mathbf{\Theta}_{\mathrm{m}}^{\mathrm{aft}}, \mathbf{\Theta}_{\mathrm{a}}^{\mathrm{aft}}), \forall t_i \in \mathbf{t}_{\mathrm{m}}^{\mathrm{aft}}. \tag{40}$$

where $S_{\mathrm{Total}}^{\mathrm{aft}}$ is given in Equation 35.

Cells born after $T_{\mathrm{d}}$ and undergo apoptosis after $T_{\mathrm{d}}$:

$$L_5(\mathbf{\Theta}_{\mathrm{m}}^{\mathrm{aft}}, \mathbf{\Theta}_{\mathrm{a}}^{\mathrm{aft}}, T_{\mathrm{delay}}, q; \mathbf{t}_{\mathrm{a}}^{\mathrm{aft}}) = \prod_{t_i} (1-q) h(t_i; \mathbf{\Theta}_{\mathrm{a}}^{\mathrm{aft}}) S_{\mathrm{Total}}^{\mathrm{aft}}(t_i; \mathbf{\Theta}_{\mathrm{m}}^{\mathrm{aft}}, \mathbf{\Theta}_{\mathrm{a}}^{\mathrm{aft}}), \forall t_i \in \mathbf{t}_{\mathrm{a}}^{\mathrm{aft}}. \tag{41}$$

Cells born after $T_{\mathrm{d}}$ and undergo neither mitosis nor apoptosis after $T_{\mathrm{d}}$:

$$L_6(\mathbf{\Theta}_{\mathrm{m}}^{\mathrm{aft}}, \mathbf{\Theta}_{\mathrm{a}}^{\mathrm{aft}}, T_{\mathrm{delay}}, q; \mathbf{t}_{\mathrm{stay}}^{\mathrm{aft}}) = \prod_{t_i} (q + (1-q) S_{\mathrm{Total}}^{\mathrm{aft}}(t_i; \mathbf{\Theta}_{\mathrm{m}}^{\mathrm{aft}}, \mathbf{\Theta}_{\mathrm{a}}^{\mathrm{aft}})), \forall t_i \in \mathbf{t}_{\mathrm{stay}}^{\mathrm{aft}}. \tag{42}$$

in all the above likelihoods, $t$ incorporates the $T_{\text{delay}}$ while calculating the hazard rate and the survival probability.

The full combined Likelihood is a product of all the 6 likelihoods

$$L_{\text{full}}(\mathbf{\Theta}_{\text{m}}^{\text{aft}}, \mathbf{\Theta}_{\text{a}}^{\text{aft}}, T_{\text{delay}}, q|\mathbf{t}) = \prod_{i=1}^{6} L_i(\mathbf{\Theta}_{\text{m}}^{\text{aft}}, \mathbf{\Theta}_{\text{a}}^{\text{aft}}, T_{\text{delay}}, q; \mathbf{t}_{\text{i}}) \quad (43)$$

| Conc. | $\mu_m$ | $\sigma_m$ | $\lambda_m$ | $\mu_a$ | $\sigma_a$ | $\lambda_a$ |
|---|---|---|---|---|---|---|
| High | 22.45 | 1.67 | 0.010 | 54.26 | 13.43 | 1.52 |
| Medium | 23.74 | 2.15 | 0.026 | 53.01 | 12.0 | 0.38 |
| Low | 21.82 | 0.73 | 0.04 | 60.11 | 14.51 | 1.65 |

Table 4: Parameters of the MCMC-inferred post-cisplatin distributions for U2OS cells.

After running Metropolis Hastings MCMC using the full Likelihood, the marginal distributions of each parater were inferred, and their means reported in Table 4. The $T_{\text{delay}}$ parameter was found to be less than 3 mins. The parameters of the MCMC-inferred underlying distributions for HCT116 cells were taken from [1]. The parameter values are provided below in Table 5.

| $\mu_{birth}$ | $\sigma_{birth}$ | $\lambda_{birth}$ | $\mu_{death}$ | $\sigma_{death}$ | $\lambda_{death}$ |
|---|---|---|---|---|---|
| 27.7 | 1.5 | 0.014 | 58 | 29 | 0.5 |

Table 5: Parameters of the MCMC-inferred post-cisplatin distributions for HCT116

### 9 Models of cell fate decisions

Exponential growth-like population models (for example Model *M0*) in the main text assume that cell fates are decided by stochastic competition between single cell times to division and death after drug administration. Deviations in the estimates of decay rates from *M0* suggested that the cell fates are likely decided largely independent of the single cell times, and the simplest model for such a decision-making process would be a "biased coin-toss model" (Model *M2*). Here the fate of each cell is independently assigned at the time of drug administration with probabilities for division, death or survival. Depending on the drug concentration, the probabilities would be biased towards division and survival (lower concentration) or death (higher concentration).

While this model could successfully capture the observed decay rates in the HCT116 dataset, it failed to generate any correlations between the fates of cells closely related by lineage as measured by us and others in previous studies [7, 8, 9, 1]. Using time lapse microscopy, up to second cousins in HCT116 cells were shown to exhibit similarity in end-fate that cannot be explained by chance. Additionally, sister cells shared the same fate about 80% of the time, irrespective of whether they were born before or after cisplatin treatment [1]. These observations cannot be recapitulated by *M2*, suggesting that the biased coin-toss model is also not a sufficient description of cell fate decisions.

However, the observation of lineage correlations in cell fate do provide a guide for the basic ingredients required to build the most suitable model: (1) distinct cellular states must exist well before drug treatment, (2) the states must be inherited across at least 2-3 cellular generations, (3) the pre-existing states predispose cells to their eventual fate after drug treatment, and (4) since the fate correlations are not 100%, occasional switching between the states must occur at a rate slower than the division rate. We therefore next developed Model *M1* that incorporates all the above 4 features. Since in experiments the cells exhibit two distinct and easily distinguishable end fates on cisplatin treatment – death or survival, the simplest coarse-grained description of cell-state heterogeneity would assume the existence of two states before drug addition, a sensitive state that primes the cell to die ($S'$) and the other, a drug-tolerant state that primes the cell to survive (pre-DTP state $P'$). We modeled switching between these two states as a Markov process using a transition probability matrix $T$. Model *M1* therefore allows for the development of a sub-population of $P'$ cells by the time of drug addition, even upon starting with a purely $S'$ population. Finally, at the time of drug administration, cells in the $P'$ state either divide or stay alive while cells in $D$ state were set to die. The waiting times to their fates for cells in either of the states were sampled from the measured post-drug IMT and AT distributions respectively. Interestingly, while *M1* could explain both the decay rates and lineage correlations for any given drug concentration data, a conceptual problem arose while attempting to explain data across all the drug concentrations in the U2OS dataset. Transitions between the cell states prior to drug administration should be an inherent property of the cell type, set by the underlying dynamics of the non-genetic processes governing the state switching. Hence the transition probability matrix $T$ must be fixed across the experiments with varying cisplatin concentration, for a given cell type. Consequently, at the time of drug administration similar fractions of pre-DTP and $D$ state cells should exist across the different experiments, thereby precluding an explanation for the different decay rates observed for the low, medium, and high cisplatin datasets for U2OS cells.

Therefore we developed the last and final model (Model *M3*), which was essentially a combination of models *M1* and *M2*. Like *M1*, here too the cells could switch between 2 states $S'$ and $P'$ before drug administration. At the time of drug addition, fates were assigned to each single cell via a fate matrix $F$ (similar to *M2*). However, unlike *M2*, the fate probabilities were conditioned on the state of the cells at the time of drug administration.Therefore at time of drug administration if an extant cell was in the $S'$ state, then it had a larger chance of dying in response to the drug. Model *M3* could quantitatively recapitulate both the lineage correlations as well as the concentration-dependent population decay rates. Details of the model implementation and final parameter values of *M3* are provided in the next section.

### 10 Simulation of Model *M3* and obtaining the best parameters.

To simulate Model *M3* we chose a sequence of transition rate values and fate matrix values and ran 100 iterates of simulation for each combination of transition and fate matrix so generated. The range of transition rates are shown in Table 6. The simulations performed were for discrete state continuous time Markov processes where the states of cells were updated after every 3 minutes. The fate matrix was chosen asymmetrically based on the states of the cells. Cells belonging to the DTP state were less prone to die. Sensitive cell state on the contrary was given higher probabilities to die.

Model *M3* resulted in the closest match between simulated and experimental results, for the first decay rate post cisplatin and the lineage correlations. 100 simulations of the model were performed for each transition matrix ($T$) and fate matrix ($F$) combination in the model. For each combination of $T$ and $F$, the Euclidean distance between simulated and experimental data was constructed using the the first decay rate and lineage correlations averaged over the 100 simulations. The transition rates were first chosen from a coarse range of $1 \times 10^{-3}h^{-1}$ to $20 \times 10^{-3}h^{-1}$. After that, a fine range was kept from $9.0 \times 10^{-3}h^{-1}$ to $14.0 \times 10^{-3}h^{-1}$ that best matched the results of the experiment.

| Parameter | Lower bound $h^{-1}$ | Upper bound $h^{-1}$ |
|---|---|---|
| $\alpha$ | $9.0 \times 10^{-3}$ | $14 \times 10^{-3}$ |
| $\beta$ | $9.0 \times 10^{-3}$ | $14.0 \times 10^{-3}$ |

Table 6: The transition rate ranges shown for Sensitive to pre-DTP ($\alpha$) and pre-DTP to Sensitive ($\beta$).

The range of Fate matrix and Transition rate matrix were decided based the states of the cells. In the beginning, a coarse search for the best fitting parameters was done. The fate probabilities were specifically chosen asymmetrical for the two states. State 'S' were less prone to divide and more prone to die. Thus, state 'S' cells were assigned the following fate probabilities: $P(\text{die}|S) \in [0.5, 1)$ and $P(\text{div}|S) \in [0.0, 0.5)$. $P(\text{stay}|S)$ was calculated by using the law of total probability i.e $\sum_{\text{fate}} P(\text{fate}|S) = 1$.

A finer range was then chosen after isolating instances where the simulation results matched with those in the experiment. The finer fate matrix values are given in Table 7:

| Fate | State | lower bound | upper bound |
|---|---|---|---|
| Death | S | 0.8 | 0.95 |
| Division | S | 0.0 | 0.15 |
| Stay | S | 0.0 | 0.15 |
| Death | P | 0.05 | 0.15 |
| Division | P | 0.0 | 0.3 |
| Stay | P | 0.65 | 0.95 |

Table 7: The Fate matrix shows the probabilities for cells in states 'S' and 'P' to succumb to fates of 'Death', 'Division' or 'Stay'.

The final parameter values that gave the simulation results that best matched with those in the experiments are provided below. Transition rates used in the simulation were symmetric for transition between the two states, $9 \times 10^{-3}$ per hour. The fate probabilities for cells to die, divide and stay alive for the two states were: Sensitive state S' = (0.9125, 0.05, 0.0375); pre-DTP state P' = (0.35, 0.1, 0.55).

### 11 Barcoding simulations

For the barcoding simulations, we used Model *M3* with the final parameters obtained by matching the lineage correlations and decay rates measured in the HCT116 experiments (parameter values provided in Table 7). To mimic barcodes, we provide lineage numbers to the initial ancestor cells that the simulations of Model *M3* are started with. These lineage numbers are then maintained across cell divisions. Starting with 65 cells (lineages 1-65), the cells were expanded to approximately 400-500 cells, followed by cisplatin addition. Cisplatin addition was modeled exactly the same way as in Model *M3*, with the Fate matrix $F$ and the post-cisplatin IMT and AT distributions. Three types of barcoding simulations were performed, which are slight variants of Model *M3* – 'Control', 'NoTrans' and 'SimExpt'. In 'Control', the simulation has no cell death during the cisplatin exposure period. In 'NoTrans', the states assigned to cells at the start of the simulation are not allowed to transition, but addition of the drug is modeled exactly as *M3*. Finally, 'SimExpt' is identical to Model *M3*.

### 12 Calculation of Barcode abundance and diversity

To calculate the barcode abundance, 100 iterations each of 'Control', 'NoTrans' and 'SimExpt' variants were performed. The frequencies of each of the 65 barcodes were extracted just before cisplatin addition and after 72 hours of cisplatin exposure. Average number of cells carrying each barcode computed over the 100 iterations was defined as 'Abundance' of the barcode.

Let $n_i$ be the number of times the $i^{th}$ barcode is observed. Then the probability $p_i$ of observing the barcode is given as

$$p_i = \frac{n_i}{\sum_i n_i} \tag{44}$$

The Shannon Diversity Index (SDI), which gives a quantitative measure of barcode diversity, was then calculated as

$$\text{SDI} = -\sum_i p_i \log(p_i) \tag{45}$$

The difference in the SDI before and after drug then provides the change in diversity of barcodes.

### 13 Luria-Delbruck Analysis

Here we count the number of persisters arising in each lineage tree. The distribution of the persister count across lineage trees suggests whether or not the transition to persistence was a result of the action of the drug or if the fate of these cells was decided before drug addition. To perform Luria-Delbruck analysis on the HCT116 dataset, we extracted the number of persisters arising from each lineage tree. Using bootstrap, we randomly sampled persister cell counts from 52 out of 65 lineages, 100 times with replacement. The persister counts across 52 lineages in each of the 100 bootstrapped samples were used to calculate 100 Indices of Dispersion (IDs) using the equation below:

$$\text{ID}_i = \frac{Var[\mathbf{x}_i]}{E[\mathbf{x}_i]}, \tag{46}$$

where $i$ goes from 1 to 100, $\mathbf{x}_i$ gives the 52 persister counts in the $i$th bootstrap, $Var$ and $E$ are the variance and expectation values of persister counts in each bootstrapped sample. Using the 100 bootstrap samples, we generated a distribution of IDs which is shown in the Main text Figure 5d under 'Expt'. Similarly we performed the above calculation of IDs for model *M3* with the transition rates between the two states, $9 \times 10^{-3}$ h and fates matrix probabilities Sensitive state S' = (0.9125, 0.05, 0.0375); pre-DTP state P' = (0.35, 0.1, 0.55). To compare the simulation results against the experimentally generated ID distribution. For control, we ran simulations of model *M2*. Here, the fates of cells were decided at the time of drug exposure independent of cell states. This variant of simulation reproduced the condition where onset of persistence occurs entirely after drug exposure. Then the bootstrap procedure mentioned above was used to obtain the ID of this Control.
